# Rapid Floral and Pitcher Scent Diversification in Carnivorous Pitcher Plants (Sarraceniaceae)

**DOI:** 10.1101/079947

**Authors:** Winnie W. Ho, J. Nathan Kutz, Julienne Ng, Jeffrey A. Riffell

## Abstract

Plant volatiles play vital roles in signaling with their insect associates. Empirical studies show that both pollinators and herbivores exert strong selective pressures on plant phenotypes. While studies often evoke the assumption that volatiles from floral and vegetative tissues are distinct due to strong pollinator-mediated selection operating on the flowers or selection from herbivores acting on the leaves, explicit tests of these assumptions are often lacking. In this study, we examined the evolution of floral and vegetative volatiles in the North American (NA) pitcher plants (Sarraceniaceae). In these taxa, insects are attracted for both pollination and prey capture, providing an ideal opportunity to understand the evolution of scent compounds across different plant organs. We collected a comprehensive dataset of floral and vegetative volatiles from across the NA Sarraceniaceae. We used multivariate analysis methods to examine whether volatile profiles are distinct between plant tissues, and investigated rates of scent evolution in these unique taxa. Our major findings revealed that (i) flowers and pitchers produced highly distinct scent profiles, consistent with the hypothesis that volatiles alleviate trade-offs due to incidental pollinator consumption; (ii) across species, floral scent separated into distinct regions of scent space, while pitchers showed little evidence of clustering – this may be due to convergence on a generalist strategy for insect capture; and (iii) rates of scent evolution depended on tissue type, suggesting that pollinators and herbivores differentially influence the evolution of chemical traits. We emphasize the need for additional functional studies to further distinguish between volatile functions.

## INTRODUCTION

Volatile organic compounds emitted from plants play critical ecological roles in community structure, ecosystem function (Andrews et al. 2007; Kessler et al. 2011; Pichersky and Gershenzon 2002), and vital evolutionary roles in the form of reproductive isolation and speciation (Amrad et al. 2016; Bischoff et al. 2015). Although emitted as complex mixtures composed of tens to hundreds of compounds, certain constituents in the mixture operate as critical “signals” to mediate these ecological and evolutionary processes (Schäffler et al. 2015; Terry et al. 2007). Surprisingly, and in contrast to literature on insect pheromones (Ma and Roelofs 2002) (Lassance et al. 2010), little is known about the specific evolutionary forces acting on plant scent constituents (volatiles) that operate as signals (but see (Schiestl 2010; Weber et al. 2016), even though there is abundant empirical evidence showing that scent is an important part of many angiosperm phenotypes. For example, floral traits are often subject to strong sexual selection (Queller 1983), with volatiles as likely targets of selection from pollinators (Ashman 2009; Delph and Ashman 2006; Waelti et al. 2009). Natural selection in association with herbivores also play important roles (Agrawal et al. 2012; Carmona and Fornoni 2013), and given the importance of volatiles in mediating plant-pollinator (Knudsen and Tollsten 1993; Raguso 2008) and -herbivore interactions (De Moraes et al. 1998; Kessler and Baldwin 2001; Paré and Tumlinson 1999; Sheridan and Karowe 2000), we should expect to see differential signatures of pollinator-mediated selection on floral function and natural selection on pitcher leaf function, acting on scents emitted by flowers and vegetation. Yet despite this strong expectation that plant volatile traits should evolve in different ways between flowers and leaves, these predictions have not yet been explicitly tested.

Carnivorous plants are especially useful for teasing apart how volatiles might evolve across tissues. In the North American (NA) *Sarracenia*, modified leaves form conical pitchers that trap a wide variety of insect types, including pollinators, for supplemental nutrition (Ellison and Gotelli 2001; Jürgens et al. 2009; Jurgens et al. 2012). In at least several species of *Sarracenia*, pitchers produce detectable levels of volatiles of diverse volatiles, including oxygenated aromatics (e.g., methyl benzoate, 2-phenyl ethanol), and monoterpenes (e.g., (*Z*)-β-ocimene, limonene), that can function as attractants to many insect pollinators (Jürgens et al. 2009). Similarly, in the pitcher plant *Nepenthes rafflesiana*, the upper pitchers emit a complex scent composed of volatiles that are attractive to both insect prey and pollinators, including aromatics (e.g., methyl benzoate, benzyl alcohol), monoterpenes (e.g., (*E*)-β-ocimene, linalool, and limonene) (Di Giusto et al. 2010). Because both flowers and leaves emit scent, the capture of pollinators can occur, leading to a conflict between plant reproductive and nutritive functions. Several studies have shown that visual cues, as well as spatial and temporal separation of pitchers and flowers, may help alleviate this potential conflict (reviewed by Jurgens et al. 2012). While pollinators and prey are the most obvious players in carnivorous plant taxa, pitcher plants are also subject to intense herbivory (Moon et al. 2008; Stephens and Folkerts 2012). Carnivorous plants may thus be under different selective pressures to alleviate pollinator-prey conflict and herbivory, and this may be reflected in the scent composition of the pitchers and flowers. To date, few studies have simultaneously analyzed floral and vegetative scent composition and evolution within a phylogenetic context.

The North American (NA) carnivorous pitcher plants (Sarraceniaceae) are an excellent model to examine how volatiles may be shaped by different selective pressures. Flowers are typically pollinated by bumblebees (or in smaller species, solitary bees), appearing briefly in the spring and emit high-intensity (to human) scents (Folkerts et al. 1999; Meindl and Mesler 2011; Schnell 1983). In contrast, leaf tissues can persist for months and represent a long-term investment throughout the growing season. These can be subject to intense vegetative damage by endemic noctuid moths, a primary herbivore of *Sarracenia* spp. (Moon et al. 2008; Stephens and Folkerts 2012). In this study, we combined a phylogenetically comprehensive sample of NA Sarraceniaceae flower and pitcher volatile data with multidimensional data analysis techniques to examine two major roles of pitcher plant scent in mediating insect attraction: (1) To determine if pitcher scent represents convergence on an attractive volatile phenotype (e.g. floral mimicry) (e.g. Schaefer 2009, Di Giusto 2010), or if pitcher scent has diverged from floral scent; and (2) To establish the potential functional roles responsible for floral and vegetative scent divergence by examining rates of scent evolution in compounds known to be associated with bee-attraction, and by examining the rates of scent evolution of compounds known to be associated with plant herbivory. Our data show strong scent divergence between flowers and pitchers, but similar profiles in pitcher scent across species. Furthermore, rates of compound evolution in pitchers and flowers suggest conflicting selective pressures acting on pitchers and flowers.

## METHODS AND MATERIALS

### Plant material and volatile sampling

We conducted a comprehensive sampling of the NA Sarraceniaceae, which included all major recognized species complexes (Mellichamp 2009; Stephens et al. 2015), in addition to one hybrid South American species. Plants were washed, bare-rooted, and vegetation removed prior to potting in a 40:60 mix of pumice and peat moss. Pots were kept outdoors (Seattle, WA 47.606° N, 122.332° W) in an artificial bog and bottom watered using the municipal water supply (unfertilized). Plants were sourced primarily from the University of Washington botany greenhouse collections and supplemented with accessions from commercial suppliers (S1-B). Approximately half the clones of *S. flava*, *S. rubra gulfensis*, *S. oreophila*, and *S. purpurea* were raised in a climate-controlled growth chamber 25-30°C, 16:8 light:dark to ensure that floral production was initiated before being transferred outdoors. There were no obvious effects on scent clustering for this group of plants (S2-C). Volatiles were collected using established plant headspace collection methods (Byers et al. 2014; Raguso and Pellmyr 1998; Riffell et al. 2008) from flowers during anthesis, and from mature pitchers (terminal edge fully opened for < 14 days) covered with pollination bags (1mm mesh) to prevent the incursion of macroscopic insects (NA species: n_(flower)_ = 4-20, n_(pitcher)_ = 6-22; n_(total)_ = 358 samples). Briefly, plants were enclosed for 24h to control for circadian variation, using Nylon bags (1.5L, Reynolds; IL, USA), and scented headspace air pulled through cartridges containing 50mg of PorapakQ adsorbent (mesh size 80-100, Waters Corp.; MA, USA). Empty nylon bags were run in parallel with all plant samples and were subtracted to control for ambient environmental contaminants. Traps were eluted in 600μL of 98% HPLC grade hexane, and headspace extracts were stored at -80°C in 2mL glass vials with Teflon-lined caps prior to analysis. 3μL of each extract was injected into and analyzed on an Agilent 7890A gas chromatograph coupled to a 5975C Network Mass Selective Detector (Agilent Technologies; Palo Alto, CA, USA). Samples were run on a DB-5 GC column (Agilent; CA, USA; 30mx250μmx0.25μm) with a constant 1.2mL/min flow of helium carrier gas. Initial oven temperature was set at 45°C for 4 min, followed by a heating ramp of 10°C/min to 250°C, with a final 8min isothermal hold.

In addition to dynamic sorption methods, headspace samples were confirmed using solid phase microextraction fibers were taken using 75μm Carboxen-Polydimethylsiloxane (CAR-PDMS) fibers (Supelco; PA, USA). Plants were similarly enclosed in Nylon collection bags, and the SPME holder needle was punctured into the bag prior to exposing the fiber at a distance of 1cm under the flower or between the peristome edges of pitchers. Samples were taken for 1h prior to desorption on the GCMS (conditions as above). Controls for both SPME and dynamic sorption samples were taken concurrently using empty Nylon bags for all samples. Volatile collections were robust to sampling method, and both CAR-PDMS SPME fibers and dynamic headspace sampling recovered similar clustering during our SVD analysis (S2-C). For consistency, all phylogenetic analyses and any calculations of emission rates were conducted only on data from dynamic headspace samples.

Chromatogram peaks were tentatively identified using the NIST08 mass spectral library (discarding compounds with a <30% match), and verified using available authentic standards and published Kovats indices in combination with retention indices calculated from C7-C30 alkane standards. A more conservative threshold was used for the dynamic headspace samples in the phylogenetic analyses: we included compounds only if they had a library match of >70%. This process was semi-automated using modified Python 2.7 scripts customized for data analysis in our lab (Clifford 2017). Chemical abundance was calculated by integrating mass spectral peaks from total ion chromatograms (ChemStation, Agilent Technologies). Emission rates are presented as ng/g wet mass/24h (Table 1), calibrated using linear fits of dilution series taken from caryophyllene and limonene standards, both widespread compounds found in *Sarracenia* spp., as well as from a toluene standard. Because the extent of ionization can vary even within chemical classes, emission rates were based on an average of all three standards. We sorted through an initial matrix of 299 putatively identified volatiles in order to isolate compounds that best explained divergence and correlation across species and tissues. Volatiles with no variance across species were removed from the SVD dataset, as they are not useful for characterizing differences across species or individual plants (but are provided in Table 1).

### Data processing for plant scents

Plant volatile traits can comprise hundreds of individual compounds; we thus performed a model reduction to retrieve volatile features most important for explaining variation across species and tissues. The singular value decomposition (SVD) is a statistical procedure that provides a data decomposition that separates variables into a set of orthogonal modes. These modes can then optimally capture linearly uncorrelated, orthogonal axes that extract the maximal variance across a data matrix. The SVD is the underlying algorithm used in principal component analysis (PCA); unlike PCA however, the SVD does not require that the data for each plant have mean-zero and unit variance. The measured intensity levels of individual components can vary over many orders of magnitude (up to 10^9^ arbitrary MSD units), with reasonable intensity levels being greater than 10^4^. Peaks with intensity levels two orders of magnitude below this could not be distinguished from background noise fluctuations in the measurements. We therefore used noise-reduction thresholds in combination with SVD to extract a sparse, but representative matrix of correlated chemical representations of floral and vegetative profiles. Raw data matrices were log transformed so that compounds with the strongest intensity (on the order of 10^9^) did not render the remaining, but significant, data irrelevant (e.g. in the range of 10^4^ to 10^7^). Because each plant has only a positive intensity level of only a small subset of the compounds measured, the data is sparse (mostly zeroes) in the space of possible volatiles, making it inappropriate to mean subtract (as in PCA) and render the majority of volatiles zero and non-negative. Instead, we used the SVD to extract the dominant correlated expression levels of volatile production. The SVD modes extract the most meaningful complex volatile bouquets derived from the hundreds of individual compounds. Final scent distances across species were calculated as the distance between centroids across the three main axes of scent divergence, SVD modes 2-4. Using a similar procedure, we also conducted nested SVDs to examine modules of chemical divergence only across flowers, and only across pitchers. The first SVD mode is not highly informative since it represents the average chemical profile of the entire dataset. All clustering analyses were performed in Matlab R2016a (see S2-A for analysis workflow).

### Constructing a time-calibrated phylogeny

As the most recent *Sarracenia* phylogeny (Stephens et al. 2015) was not time-calibrated, we used the same data to generate a time-calibrated molecular phylogeny (Fig 1). First, we imported Stephens et al.’s (2015) 199 nuclear gene alignment into BEAST 2 (v. 2.4.0; Bouckaert et al. 2014) and constrained the tree to ensure the resultant tree had the same topology as Stephens et al.’s (2015) species tree. We calibrated the tree following divergence time estimates from a family-level phylogeny using a normal prior distribution (Ellison et al. 2012): the *Sarracenia* crown node was constrained to a mean of 4.18 Ma (offset 2.0, sigma 1.5), the *Sarracenia* stem node was constrained to a mean of 22.76 Ma (offset 14.0, sigma 5.0), and the stem node of *Darlingtonia, Heliamphora* and *Sarracenia* was constrained to a mean of 34.91 (offset 25.0, sigma 5.0). We conducted two runs of 50 million generations, sampling every 5000 generations, and used Tracer v1.6 (Rambaut et al. 2015) to verify that both runs reached stationarity and converged on the posterior distributions of trees. We discarded 10% of the trees as burn-in, as identified in Tracer, and then combined and summarized trees as a maximum clade credibility (MCC) tree using LogCombiner v. 2.4.0 and TreeAnnotator v. 2.4.0 (both programs are provided as part of the BEAST package). All nodes were highly supported (posterior probabilities=1). Prior to analyses, we pruned species not included in the study from the MCC tree. As Stephens et al. (2015) sampled a number of individuals per species, we chose one representative individual to represent a species and pruned all other individuals from the MCC tree prior to analyses. As individuals from most species formed a monophyletic group, pruning taxa from the same clade will not change branch lengths for the species or affect results. All analyses of scent evolution (see below) were conducted on the MCC tree as well as 100 trees sampled from the posterior distribution of trees from BEAST to incorporate branch length uncertainty. From these 100 trees, we extracted the 95% highest posterior density credibility interval (CI). Additionally, as the placement of *S. rubra ssp. gulfensis* was unresolved in Stephens et al.’s (2015) species tree, we conducted all analyses with *S. rubra ssp. gulfensis* as sister to *S. jonesii*, as sister to *S. alata*, and sister to both *S. jonesii* and *S. alata*. As the results were primarily qualitatively similar, unless mentioned, we here only report the results conducted on the tree where *S. rubra ssp. gulfensis* is sister to both *S. jonesii* and *S. alata*.

**Figure 1.**
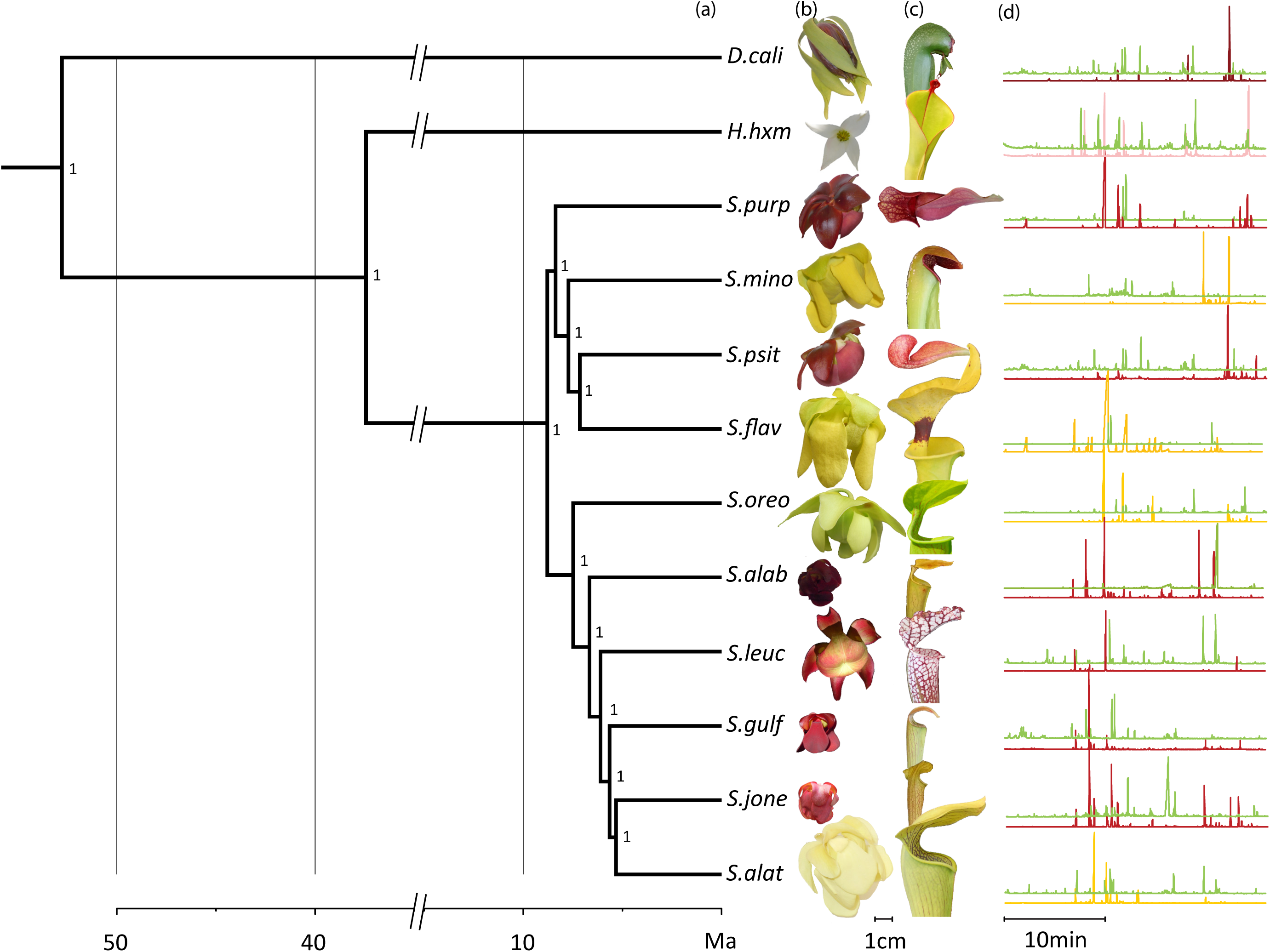
MCC tree from the BEAST analysis of the (a) Sarraceniaceae with (b) flowers, (c) pitchers, and (d) scent chromatograms (upper green trace = pitcher; lower trace = flower); amplitudes are scaled for ease of visualization (*H.hxm* flower photo courtesy of the UW botany greenhouse). Numbers on nodes represent posterior probability values.

### Analysis of scent evolution

To examine rates of volatile evolution across flowers and pitchers, we assigned compounds from the volatile dataset as being involved in possible pollinator-mediated interactions (pollinator ‘P’; four compounds), or as involved in herbivory-related interactions (‘H’; seven compounds). Because we do not yet have specific functional data on how volatiles mediate plant-insect interactions in the Sarraceniaceae and individual compounds may be associated with multiple functions, we based these designations on published data from typical pollination syndromes (*Bombus* spp. or solitary bees, which are the primary floral visitors in the NA Sarracaeniaceae) (Folkerts et al. 1999; Meindl and Mesler 2011; Schnell 1983) for pollinator and insect attraction function, and defense-related volatiles for herbivore related function. Furthermore, an additional criterion was that these volatiles elicited responses in receiver behavior and olfactory physiology (Table 1). Finally, we emphasize that this analysis requires that volatiles be produced in both the flowers and pitchers of multiple species, in order to directly compare diversification across each tissue. We therefore pruned this dataset in order to include volatiles that were produced in the flowers or pitchers of at least four species each (range=4-13).

Recent development of multivariate comparative methods now allows analysis of high-dimensional multivariate phenotypes in a phylogenetic context (e.g. Adams and Otárola-Castillo 2013; Clavel et al. 2015). While these methods were developed and have typically been used for morphological shape data, we here apply these methods to scent data; another high-dimensional, multivariate trait. We estimated phylogenetic signal in pitcher and flower scent to test whether phylogenetic relatedness influenced scent using K_mult_ (Adams 2014a), a multivariate generalization of Blomberg’s K statistic (Blomberg et al. 2003). We tested whether scent abundance (volatile emissions analyzed as a continuous variable) and/or scent composition (as measured by absence/presence of volatiles) exhibit phylogenetic signal. We used 1000 permutations to determine whether phylogenetic signal is significant compared to that expected under a Brownian motion model of evolution.

To test whether pitcher and flower scent evolved independently in the NA pitcher plants, we first evaluated whether pitcher and flower scent abundance and/or composition co-varied with one another across the phylogeny, following Adams and Felice (2014). To test the significance of the correlation, we permuted the phenotypic data on the tips of the tree 1000 times, each time calculating the correlation scores to which the observed correlation score was compared. Second, we estimated the net evolutionary rate over time for the scent emitted by each organ using σ^2^_mult_ (Adams 2014b). As floral volatiles tend to be emitted at higher intensities than pitchers (see Results), we used proportional data to standardize scent across flowers and pitchers. To assess significance, the ratio between pitcher and flower scent was compared to 1000 phylogenetic simulations in which data on the tips are obtained under Brownian motion using a common evolutionary rate for all traits. We further examined the evolution of different types of scents by estimating σ^2^_mult_ for flower and pitcher volatiles that have previously been associated with bee attraction and those that have been associated with herbivory deterrence (Table 1). All analyses of scent evolution were conducted using the geomorph R package (Clavel et al. 2015).

### Chemical versus temporal and spatial divergence

To determine whether increased potential for pollinator-prey conflict was related to chemical divergence between flowers and pitchers, we conducted a linear regression using combined blooming periods and between-organ height differences. To account for shared phylogenetic history among species (Felsenstein 1985), we computed phylogenetic independent contrasts for an index combining temporal and spatial data, and for the chemical difference between flowers and pitchers using the pic function in the ape R package (Felsenstein 1985). We used the resulting values to conduct a linear regression to test whether the level of chemical divergence between flowers and pitchers was inversely related to the level of spatial and temporal divergence of these two traits.

Data on the temporal and spatial separation of flowers and pitchers were compiled from the Flora of North America (FNA) (Mellichamp 2009). This method provides a standardized way to collect phenology data, but we emphasize that future *in situ* studies will be crucial to account for local and population level variation (Stephens et al., unpubl. data). Chemical distances were calculated as linear distances between centroids from data visualized in modes 2-4 of the SVD. Temporal separation was coded based on the months provided in FNA on flowering time, and on descriptions of pitcher opening periods. Descriptions were conservatively coded as follows: “shortly after first flowers” = 0.1; “with or after flowers” = 0.5; “with or just prior to flowers” or “appearing with flowers, continually” = 1. Spatial separation distances were calculated as the percent overlap between given height ranges for flowers and height ranges for pitchers (S1-C). Indices of physical separation were calculated by scaling spatial separation of flowers and pitchers with the degree of temporal separation: that is, if there is complete temporal overlap, then physical separation depends only on the height difference between flowers and pitchers.

## RESULTS

### Scent composition was dominated by terpenoids, benzenoids, and aliphatics

Volatile emissions in the NA Sarraceniaceae were dominated by mono- and sesqui- terpenes (48% of detected compounds), and included major contributions from limonene, α-pinene, and caryophyllene. Aliphatic emissions (e.g. tridecane and pentadecane) were also widespread (34%). Finally, benzenoid emissions, including a number of aromatic esters, comprised 18% of detected volatiles (Table 1). Species-specific compounds accounted for 31% of emitted compounds, and the total number of compounds detected from each species ranged from 39 volatiles in *S. leucophylla* and *S. psittacina*, to 84 in *S. alabamensis*. Flowers tended to emit greater quantities of scent (ng/mg wet weight/h) and overall, produced a greater number of volatiles (p=0.01, t(38)=2.7). This difference in volatile number was driven by an increase in the diversity of terpenoids (p<0.0001); the number of benzenoids and aliphatics did not differ across tissues (p>0.1).

### Volatile mixtures were highly distinctive across pitchers and flowers

Species (Fig. 2a) and organs (Fig. 2b) in the NA Sarraceniaceae were readily distinguished on the basis of scent (Fig. 2c). Across species, there was marked scent divergence across flowers, while clustering was not well-defined across pitchers from different species. In the SVD space, across-species separation in pitchers volatiles was very low (Fig. 2c), and the mean spread was more than six times greater in flowers than pitchers (variance 5.2x10^-3^(flower) vs. 0.84x10^-3^ (pitcher)). Furthermore, within each species, flowers and pitchers were highly divergent (Fig. 2c).

**Figure 2.**
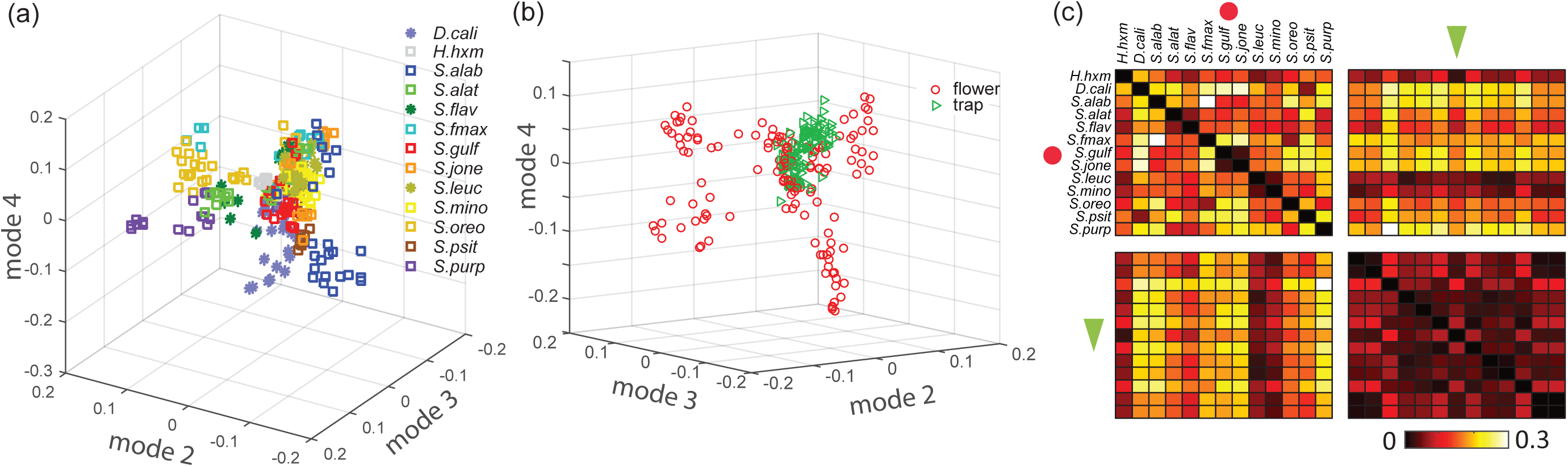
Scent differences shown (a) as scatterplots across species, (b) across tissues, and (c) as a heatmap across tissues and species. Scatterplots are adjusted to show the most important axes of separation. The color bar shows the magnitude of scent divergence: bright colors represent a high degree of separation and dark colors represent low separation. Flowers = red circles; pitchers = green triangles. Species names are abbreviated as the first generic initial, followed by 3-4 letters from the specific epithet; full names and abbreviations are provided in accession table S1-B.

### Covariance in volatile composition across NA pitcher plants

We ran further nested classifications to examine how volatiles covaried within tissues. In flowers, the SVD revealed that across-species floral composition involves strongly covaried expression of terpenoids, including caryophyllene, sabinene, *β*-pinene, and *β*-myrcene. Superimposed on this primary floral mixture, the SVD second mode reveals separation of volatiles along two directions, generating first, a module characterized by covaried production of α-curcumene, (±)-linalool, cis-*α*-bisabolene, and *α*-zingiberene, and a second contrasting strategy which flowers emitted combinations of α-ionone, eucalyptol, tetradecanal, sulcatone, and a handful of terpentine derivatives, including terpinolene and α-terpineol (Fig 3).

**Figure 3.**
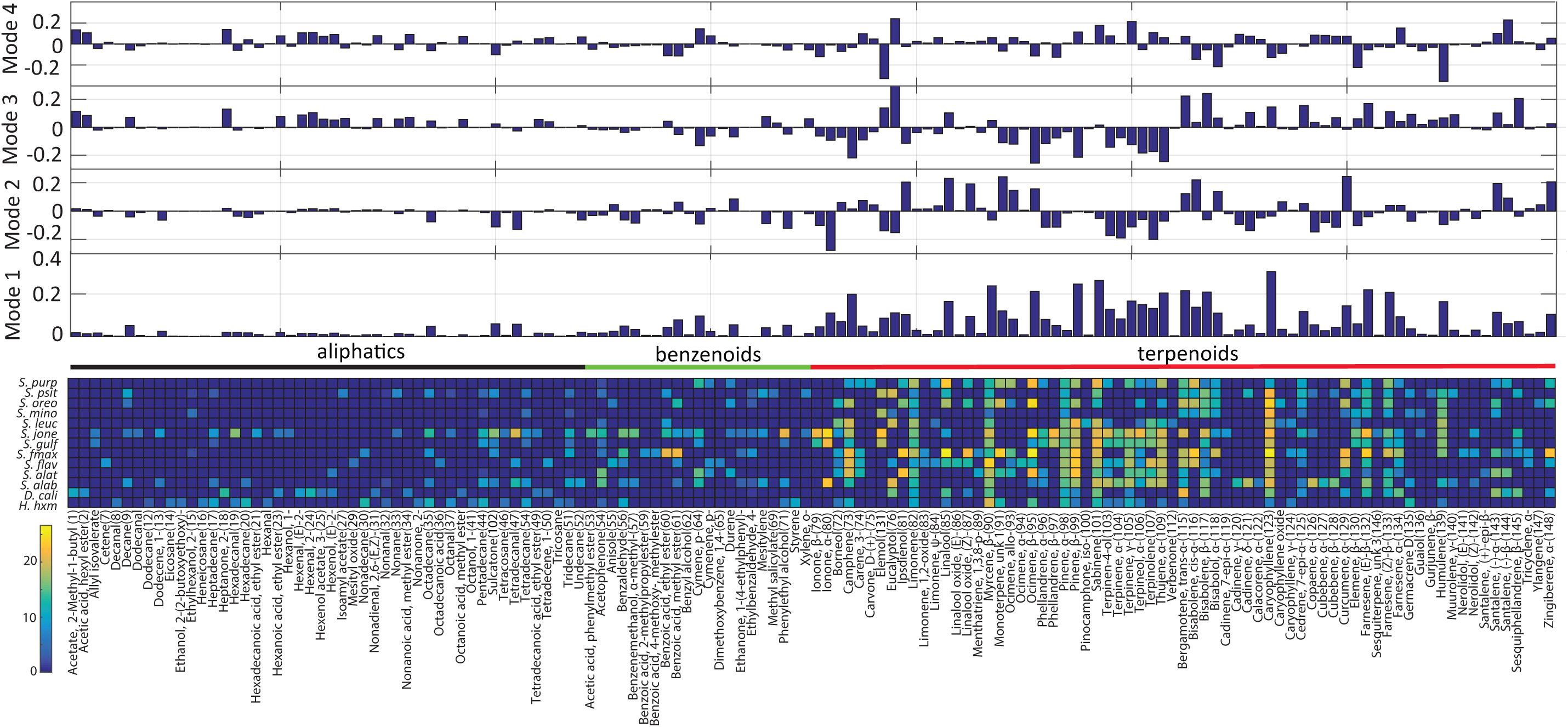
Volatiles most important for explaining floral diversity extracted from nested singular value decompositions are shown as a heatmap of relative intensity (color bar, left). Top: correlation modes 1-4.

Nested classification for volatiles within vegetative tissues found that despite low pitcher clustering in the combined organ analysis, chemical divergence across pitchers of different species is best explained by a mixture of caryophyllene, ethyl benzoate, mesitylene, *α*-farnesene, and benzaldehyde. The second SVD mode reveals two alternate pitcher strategies, the first involving predominantly terpenoid volatiles, including (*E/Z)*-*β*-ocimene, humulene, sabinene, and linalool, but also 1-octanol and (*Z*)-3-hexenol; the second involving predominantly benzenoid compounds, including ethyl benzoate, methyl benzoate, benzaldehyde, as well as 2-heptanone (S2-D).

### Phylogenetic signal and rates of scent evolution differed between flowers and pitchers

Our estimation of phylogenetic signal using continuous scent emissions data showed that floral scent did not exhibit significant phylogenetic signal (K_mult_ =1.02, [95% CI=0.75-1.07]) while pitcher scent emissions did using the MCC tree, but only some trees from the posterior distribution (K_mult_ =1.07, [95% CI=0.94-1.10], *p*<0.05). When estimating phylogenetic signal on presence/absence data, we did not find significant phylogenetic signal for either floral (K_mult_ =0.95-0.96 [95% CI=0.88-0.98], *p*=0.05) or pitcher scent (K_mult_ =0.94, *p*>0.05). Together, this indicates that the pitchers of closely related species tend to emit similar intensities of volatiles but the compounds themselves are not similar. Conversely, the flowers of closely related species do not tend to emit similar volatile bouquets or similar quantities of scent.

We found that flower and pitcher scent significantly co-varied when tests were conducted on volatile abundance (*p*<0.05) but not when composition data was used (*p*>0.05), suggesting some correlation with the overall amount of scent produced across organs, but not across compounds themselves. Further supporting that the compounds did not co-vary between organs, the net evolutionary rate for the entire set of volatiles from flowers and pitchers significantly differed, whereby floral scent evolved 30% faster than pitcher scent (σ^2^_mult.flower_ /σ^2^_mult.pitcher_= 1.3, [95% CI=1.18-1.31], *p*<0.01). When estimating the net evolutionary rate of volatiles that have been associated with bee-visitation, we found that these scents evolved 15 times as fast in flower versus pitchers (σ^2^_mult.flower.bee_ /σ^2^_mult.pitcher.bee_ = 15 [95% CI=14.23-15.48], *p*<0.01). Conversely, herbivory-related volatiles evolved greater than two times faster in pitchers than flowers (σ^2^_mult.flower.herbivory_ /σ^2^_mult.pitcher.herbivory_ = 2.4, [95% CI=2.29-2.87], *p*<0.01).

### Spatial and temporal divergence of pitcher plant scents

We found no significant relationship with the level of chemical divergence between flowers and pitchers, and the level of temporal and spatial divergence: flowers and pitchers did not produce more divergent scent bouquets, even when they matured at similar times and heights (S2-E).

## DISCUSSION

Our study revealed distinct scent divergence between flowers and pitcher leaves, consistent with the hypothesis that scent production in these tissues is subject to distinct selective pressures in the NA Sarraceniaceae. This divergence is partially explained by a greater emission of scent compounds in flowers, which emit a greater intensity and broader range of terpenoids than pitchers. Within tissues, scent variance in flowers was more than six times greater than that in pitchers (Fig 2b). Flowers are specialists and typically recruit one main pollinator, thus this disparity may result from selection for pollinator constancy amongst flowers. By contrast, the lack of distinct clustering in pitchers may reflect a more generalist trap strategy to attract a wide variety of insect genera and species. Surveys of unbagged pitchers in our study plants confirm a range of trapped insects that include dipterans (flies, mosquitoes), lepidopterans, and hymenopterans (Ho unpubl. data). Nevertheless, in some species and regions, pitchers do exhibit specialist function (e.g. Stephens et al. 2015), and understanding how volatiles interact with pitcher traits such as morphology will provide important functional data (Stephens et al., unpubl. data)

Our SVD analysis identified several suites of chemicals that covary *within* each tissue (Fig 3, S2-D). These integrated chemical modules identified within flowers and pitchers have several ramifications. First, it is widely recognized that many traits evolve in a concerted manner (Lande and Arnold 1983; Pigliucci 2003) and have the potential to constrain or facilitate evolution (Futuyma 2010; McGlothlin and Ketterson 2008). This is also the case for floral phenotypes (Conner and Sterling 1996; Smith 2016), and in wild *Brassica*, artificial selection on a single volatile compound can pleiotropically alter the scent of the entire volatile bouquet, typically by increasing emissions of non-target compounds (Zu et al. 2016). Furthermore, volatiles from similar chemical classes in the *Brassica* study were more strongly correlated than those from different classes (Zu et al. 2016). This is consistent with our analysis of pitcher compounds, and the observed benzenoid- or terpenoid-dominated strategies might represent biochemically- or genetically-constrained modules (S2-D). Nonetheless, our phylogenetic analyses revealed that *across* floral and vegetative tissues, scent did not significantly covary in emission rate or composition. This indicates that although within-tissue emissions may covary as a unit, volatiles in this taxon might be regulated independently across tissues. This concurs with other studies of floral and vegetative traits in which selection for functional independence can occur (Armbruster et al. 1999; Conner and Sterling 1996; Meng et al. 2008).

Nevertheless, a handful of floral volatiles, including limonene, caryophyllene, α-pinene, and sabinene, were also produced in pitchers. There are several possibilities for this overlap, which could result from either floral mimicry in pitchers (e.g. (Di Giusto 2010)), convergence on similar tactics for invertebrate attraction (Schaefer 2009), or, even a need for both tissues to deter herbivory (e.g. Kessler and Baldwin 2001). One intriguing possibility is that flower and pitcher scents are aligned for long distance insect attraction, and it is only at close distances that divergence is necessary to distinguish flowers and pitchers. This is consistent with a recent study showing that floral scent in *Pinguicula*, a sticky trap carnivore, attracts both pollinators and non-pollinating prey, whereas only prey are attracted to leaf scents (El-Sayed 2016). This synergistic effect of flower and leaf scent on insect attraction is also observed in other taxa (e.g. Kárpáti et al. 2013). Finally, because vegetative and floral tissues often share overlapping biochemical pathways (Kessler 2009; Berardi 2016), another possibility is that the expression levels of these compounds across flowers and pitchers are not readily decoupled.

### Rates of volatile evolution in flowers and pitchers

In many angiosperm systems, pollinators and herbivores are forceful drivers of floral and vegetative diversity. Here, we found that floral scent evolved 1.3 times faster than scent from trapping pitcher leaves, raising the possibility that sexual selection contributes to volatile diversity in the outcrossing NA *Sarracenia*. We investigated this further by examining specific volatiles, found in *both* flowers and pitchers, that were electrophysiologically and behaviorally relevant to *Bombus* and solitary bee pollinators in this clade. This uncovered an even stronger effect in which these volatiles evolved 15 times more rapidly in flowers than in pitchers. Our results reveal pollinator-mediated selection may have an outsized importance on the rates of floral scent evolution in the NA pitcher plants.

Although scent from pitcher leaves evolved more slowly than floral scent, we found that compounds associated with herbivory still evolved at more than double the rate in pitchers than in flowers. In the NA *Sarraceniaceae*, one of the chief herbivores are noctuid moths (genus: *Exyra*). Vegetative damage from these pitcher plant specialists (Stephens and Folkerts 2012) can exert strong selective pressure on pitcher traits, reducing plant size and leaf growth (Moon et al. 2008). Our data indicate herbivore-associated compounds evolved much more quickly in pitchers than in flowers, suggesting that herbivory, likely from *Exyra* damage, has played a significant role in the evolutionary history of the *Sarracenia* pitcher plants. Together, these results provide the impetus for integrative studies that will not only link scent production with specific pollinator and herbivore interactions, but which will also explore the functional consequences of these interactions on pitcher plant fitness.

### Summary and Conclusions

In carnivorous plants, insects function as both pollinators and prey. This unusual life history gives rise to the pollinator-prey conflict (Juniper et al. 1989), a trade-off which is most apparent in outcrossing, pollen-limited species (Jurgens et al. 2012) like the NA Sarraceniaceae (Meindl and Mesler 2011; Ne’eman et al. 2006; Sheridan and Karowe 2000). Flower and pitcher scents were highly distinct, consistent with the hypothesis that pitchers and flowers might target private sensory channels to alleviate pollinator-prey conflict. One important question then, is whether species with a greater potential for conflict (i.e. less physical separation between flowers and pitchers), should produce flower and pitcher scents that are more divergent. This was not the case in our study, and one possibility is that volatiles are not be involved in alleviating pollinator-prey conflict. However, because our data were standardized from published records, both (i) field data from specific populations would provide greater precision in quantifying pollinator-prey conflict, and (ii) *in situ* temporal data on floral and pitcher phenology, especially with respect to volatile emissions across different seasons (Stephens et al., unpubl. data) would be especially useful in evaluating the potential for conflict. Such data will also be useful in interpreting scent variance across individuals, populations, and species. Lastly, we also emphasize the need for functional data on the sensory systems of different insect guilds (pollinators, prey, herbivores) to distinguish between these possibilities.

There is now strong evidence that animal mutualists and antagonists can have robust effects on plant scent, and there is increasing information for how these forces influence scent evolution and volatile diversity, especially with respect to the sensory ecology of the receivers (Raguso 2008; Reisenman et al. 2010). This study reveals that the functional consequences of volatile signaling can have strong effects on rates of scent evolution, and recognizes the outsized influence of pollinator-related volatile traits on floral scent evolution. We also re-emphasize the importance of physiological studies that specifically target the olfactory sensory biology of Sarraceniaceae mutualists and antagonists, as well as data on how pollinator, herbivore, and prey interactions interact to influence plant fitness. These studies, along with longer-term selection experiments, are crucial for distinguishing whether scent modularity results from biochemical constraints, or from insect-mediated ecological selection. Finally, we suggest that while the traditional emphasis on prey capture in defining carnivorous plant phenotypes is a useful one, our framework should be expanded to include generous roles for herbivore- and pollinator-mediated selection, at least in the context of scent evolution.

## SUPPLEMENTARY MATERIAL

Source material accessions are stored at the University of Washington Herbarium (S1-B). Primary data files and resources for data analyses are available upon request.

## ACKNOWLEDGEMENTS

We are indebted to J. Milne, J. Addington, P. Beeman, MBRS, and the dedicated staff at the UW Botany Greenhouse for source material and collections maintenance. K. Aoki, S. Huang, E. Kuo, and J. Miao assisted with volatile collections; E. Roth, C. Rusche, and especially M. Clifford, were instrumental in developing code to automate our data analysis. We also thank J. Stephens and A. Kessler for helpful discussions and advice on this manuscript and project. We acknowledge the support of NSF PRFB-1401888 (WWH), NSF IOS-1354159 (JAR), the Human Frontiers in Science Program HFSP-RGP0022 (JAR), and an Endowed Professorship for Excellence in Biology (JAR).

---

Supplemental Figure S1. (A) Table of additional volatiles (B) Table of Accessions (C) Physical and chemical separation of flowers and pitchers.

Supplemental Figure S2. (A) Data workflow and dimension reduction analyses (B) Scatterplots and scree plot of scent diversity (C) Scent variation by species, tissue, and sampling method (D) SVD of pitcher scent (E) chemical vs. spatiotemporal separation.

